# Chronic exposure to predator cues fails to elicit plastic responses or transgenerational effects in banded crickets

**DOI:** 10.1101/2023.12.05.570167

**Authors:** Jeremy Dalos, Ned A. Dochtermann

## Abstract

Plasticity is a major feature of behavior and particularly important for how animals respond to predators. While animals frequently show plastic responses when directly exposed to predators, with these exposures even leading to permanent behavioral changes and transgenerational effects, whether indirect cues of predator presence can elicit similarly severe responses is unclear. We exposed banded crickets (*Gryllodes siglattus*) to cues of predator presence throughout development and compared their behavior—as well as the behavior of their offspring—to individuals who had not been reared in the presence of predator cues. Contrary to findings in both *G. sigilattus* and related species, we did not detect either developmental plasticity in the form of differences between adult behavior or differences in offspring behavior. These findings suggest that chronic exposure to cues of predator presence have a substantially different affect on behaviors than does direct exposure to predators. How habituation might interact with developmental plasticity and transgenerational effects requires further investigation.

**Significance Statement:** Previous research has established that exposure to predators elicits behavioral plasticity, including life-long effects, as well as transgenerational effects. Here we show that chronic exposure to cues of predator presence throughout development, with a resulting potential for habituation, results in neither differences in adult behavior or transgenerational effects. This suggests an important role for habituation in how plasticity manifests within and between generations

## Introduction

Plasticity is a key characteristic of behavior (Snell-Rood 2013) and responsible for at least 75% of behavioral variation on average (Dochtermann et al. 2019). Plasticity in how individuals respond to predators can be particularly important for survival as different responses may be required depending on the prey individual’s state, for different predators, or predators of a single species but in different states (Van Buskirk 2001, Preisser et al. 2007, Kikuchi et al. 2023).

This plasticity in response to predators is often closely related to the context of risk posed by particular predators. For example, Palmer and Packer (2021) examined the behavioral responses of three species of herbivores—blue wildebeest (*Connochaetes taurinus*), plains zebra (*Equus quagga*), and impala (*Aepyceros melampus*)—to models of predators that differed in hunting styles and predation threat. Of the predator species, spotted hyenas (*Crocuta crocuta*) and wild dogs (*Lycaon pictus*) are primarily coursing predators that preferentially preyed on impalas while cheetahs (*Acinonyx jubatus*), and lions (*Panthera leo*) are primarily ambush predators with cheetahs posing greater risk to impalas and lions posing greater risk to wildebeest and zebra (Palmer and Packer 2021). Risk specific responses to cues of predator presence, like those found by Palmer and Packer (2021), are common in arthropods (e.g. Binz et al. 2014, Miller et al. 2014), fish (e.g. Eklöv and VanKooten 2001), mammals (e.g. Makin et al. 2017), and birds (e.g. Kalb and Randler 2019).

Predator-induced plasticity can also result in the expression of transgenerational effects. These effects can be particularly important to fitness if they “prime” an individual to rapidly or dramatically respond to predators and cues of predators—i.e. anticipatory transgenerational effects (Marshall and Uller 2007). A particularly comprehensive examination of anticipatory transgenerational effects for behavior was conducted by Storm and Lima (2010) who exposed gravid field crickets (*Gryllus pennsylvanicus*) to wolf spiders (*Hogna carolinesis*) with waxed chelicerae for ten days, allowing for non-lethal encounters with predators. These encounters resulted in offspring of exposed crickets displaying freezing behavior 27% more often than offspring of unexposed crickets in subsequent behavior trials. Additional testing also showed that offspring of exposed crickets had significantly greater longevity and survival probability when exposed to *Hogna* spiders in survival assays (Storm and Lima 2010). However, whether similar transgenerational effects can be elicited by chronic exposure to indirect cues of predator presence—versus the direct interactions of Storm and Lima (2010)—is unclear.

While responses to indirect cues—e.g. chemical, visual, or auditory cues but without direct contact—of predators are often reflective of responses to the predators themselves (e.g. Royauté and Dochtermann 2017, Bucklaew and Dochtermann 2020), chronic exposure to cues of predator presence might produce different responses than either direct or acute exposure to predators or cues. Specifically, chronic exposure to cues could lead to either sensitization—an increase in responsiveness with repeated exposure to a stimulus—or habituation—a reduction in responsiveness with repeated exposure to a stimulus (Thompson 2009). Habituation might be particularly likely given a lack of corresponding reinforcement. While both outcomes represent plasticity, habituation will result in an *apparent* lack of plasticity if behavioral responses return to the same baseline as individuals who are never exposed to cues prior to measurement. Further, under habituation, transgenerational effects might not be expected if regulatory and physiological responses are dampened over time along with their associated behavioral responses.

Here we reared groups of *Gryllodes sigillatus* (banded crickets) in the presence of chemical, auditory, and visual cues of leopard geckos (*Eublepharis macularius*) from an early nymphal stage until maturity. By rearing *G. sigillatus* in the presence of cues of gecko presence we were able to examine whether chronic presentation during development resulted in differences in the behavior of adults, i.e. “developmental plasticity” (West-Eberhard 2003, Snell-Rood 2013), *and* whether this chronic exposure elicited transgenerational effects

## Materials and Methods

### F_0_ Rearing Conditions

*G. sigillatus* used in this study were from an outbred line established from individuals initially caught in California (Ivy et al. 2005) and currently maintained in Fargo, ND.

Hatchlings constituting our F_0_ population were initially reared in a single group housing container until individuals reached ∼1 cm in size. After reaching the targeted size, 150 *G. sigillatus* were moved into one of ten 37.9L terraria (15 crickets per container). Individuals were size matched based on length before being moved so that terraria contained similar sized individuals. Sex of individuals was unknown before maturity and, instead, sex and mass were determined at the time of behavioral testing (see below). There was an initial, and substantial, die-off of individuals following their being moved to terraria and throughout development (Table 1).

**Table 1.** Number of individuals by treatment and sex for the F_0_ and F_1_ generations. F_0_ unequal sex ratio was caused by unknown sex at treatment assignment.

| F<sub>0</sub> | Female | Male | Total | F<sub>1</sub> | Female | Male | Total |
| --- | --- | --- | --- | --- | --- | --- | --- |
| Control | 7 | 11 | 18 | Control | 23 | 24 | 47 |
| Exposure | 12 | 16 | 28 | Exposure | 50 | 51 | 101 |
| Total | 19 | 27 | 46 | Total | 73 | 75 | 148 |

All terraria were divided into two sections: a cricket and gecko area. The dimensions of the gecko section were 38 cm × 25 cm and the dimensions of the cricket area were 12.5 cm × 25 cm (Figure 1). In five of the terraria a mature leopard gecko was placed in the gecko section (Exposure terraria) while no gecko was present in the other terraria (Control terraria). A screen partition between the two sections extended from the floor to the roof of the terraria. This prevented both crickets and geckos from moving freely between sections.

**Figure 1.**
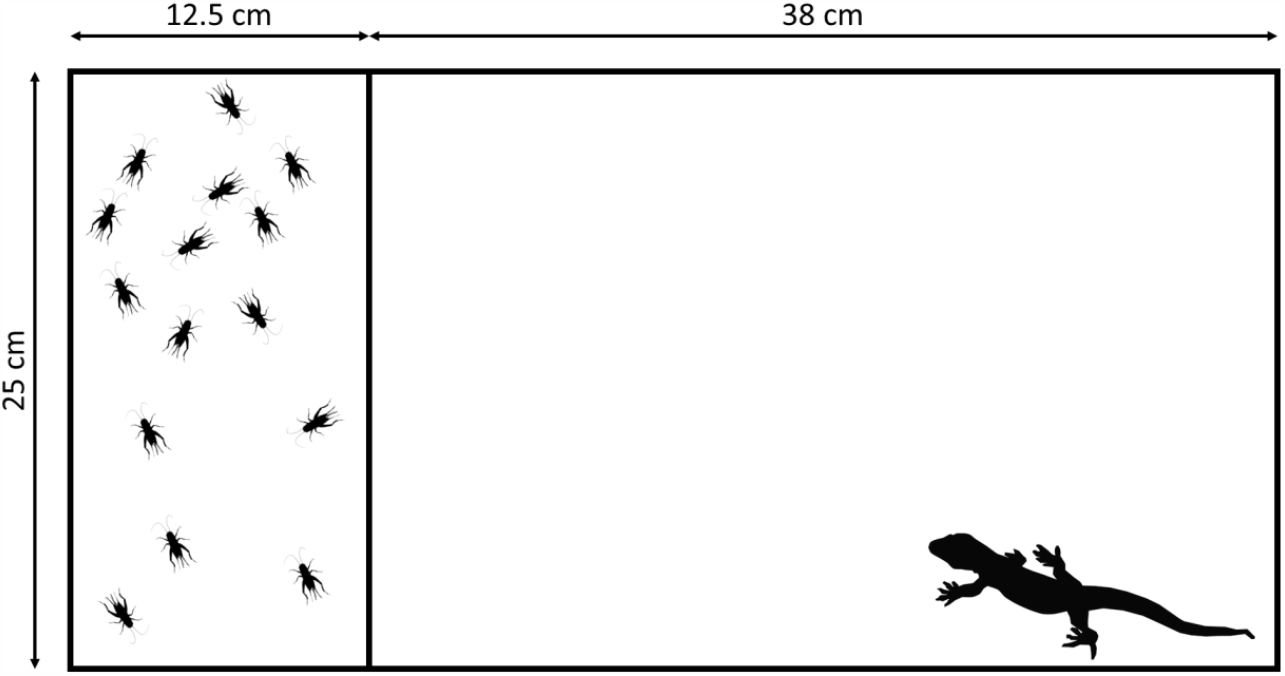
Terraria schematic used in rearing F_0_ generation. Crickets were provided with ad libitum food and water, along with shelters. Geckos were provided with water and shelter and fed live *G. sigillatus* every other day.

Each cricket portion of the terraria included a food dish, shelter (cardboard egg carton sections), and water filled glass vials with cotton plugs. Crickets were provided with *ad libitum* food (commercially purchased chicken feed following Royauté et al. (2015)). All terraria were kept under a 12:12 light:dark photoperiod with an average temperature of 29°C during the course of cricket development.

Geckos were provided with a shelter, ad libitum water, and provided 3 *G. sigillatus* as food every other day. All leopard geckos used in this study were housed and cared for according to the standards of the Animal Behavior Society (ASAB/ABS Animal Care Committee 2023) and under the regulation of the Institutional Animal Care and Use Committee of North Dakota State University (Protocol A14006, A17015, and A19067).

During rearing, the gecko and cricket sections were switched weekly by moving the barrier dividing sections and transferring crickets and geckos between them. Consequently, crickets in the exposure treatment experienced auditory and visual cues of predator presence throughout the week and were exposed to fresh chemical cues—due to skin shedding, excretion, etc.—every week. The partitions were moved in the same manner and crickets transferred between sections in the control terraria.

Once mature, individual crickets were isolated and moved into individual housing for 24 hours. Individual housing consisted of a 0.71-liter container with a transparent cover. Individual containers included *ad libitum* food, shelter, and glass vials. After the isolation period, all individuals were measured in two behavioral assays (see below).

### F_0_ Mating and F_1_ Rearing Conditions

After behavioral assays (see below), we established random pairs of male and female F_0_ individuals *within* the Exposure and Control treatments (i.e. exposure males were only mated with exposure females). We were not able to explore the full factorial combinations of treatments due to logistical constraints. Paired male and female crickets were housed in 5.7-liter containers with water, *ad libitum* food, and shelter for 24 hours and allowed to mate. After the mating period, we removed both individuals and placed them back into their respective individual housing.

After matings, we monitored the containers of F_0_ females weekly for hatchlings. F_1_ hatchlings were reared in the same 0.71-liter container with shelter, *ad libitum* food, and water as their mother until they reached 1 cm in size. Once individuals reached 1cm, we isolated nymphal crickets in individual housing until maturity was reached. Each individual housing container was also 0.71-liters with transparent covers that included food, shelter, and glass vials similar to the F_0_ generation. The F_1_ individuals were also reared under a 12:12 light: dark photoperiod at a temperature of 25-28°C. Once mature, we ran all individuals through the same behavioral assays as the F_0_ generation. 150 individuals F_1_ individuals in total were isolated into individual housing after reaching the targeted size of 1 cm, with minimal die off before maturation and testing (2/150, Table 1).

### Behavior Trials

We measured latency to emerge from shelter and activity in the presence of cues of predator presence in both F_0_ and F_1_ individuals. Both represent ecologically relevant behavioral tests for crickets. Because crickets retreat to burrows and cracks when startled, their latency to emerge from a shelter is a commonly used proxy for risk-taking propensity (e.g. Niemela et al. 2012, Kortet et al. 2014, Royauté et al. 2020). When in the presence of cues of predator presence, crickets are expected to either increase their activity (putatively an escape behavior) or freeze, depending on the hunting modality of the predator (Binz et al. 2014). Consistent with this, *G. sigillatus*, exhibit increased activity and decreased propensity to emerge from shelter following direct and acute exposure to leopard geckos (Bucklaew and Dochtermann 2020). Likewise, when in the presence of predator cues, related species show increased activity (Royauté and Dochtermann 2017, Dalos et al. 2022). We conducted behavioral assays of the F_0_ generation between May 2019 and July 2019 and between August 2019 and October 2019 for the F_1_ generation. Latency trials could not be blinded as individual ID—which included treatment group for the F_0_ generation—was included in video recordings and in video file names. Antipredator response trials were batch analyzed using Ethovision XT, blind to treatment.

### Latency Trials

We conducted latency tests in which individuals were transferred from their home containers to small artificial burrows (40 cm^3^) placed within a 34.6 cm × 21 cm arena. These artificial burrows were capped so that individuals could not immediately emerge. Crickets were forced to remain in the artificial burrow for two minutes for habituation, after which the cap was removed from the burrow. Crickets were then allowed six minutes and thirty seconds to emerge from the artificial burrow. All trials were digitally recorded. We recorded how long it took for an individual to emerge (in seconds). Individuals that did not emerge were given a maximum latency of 390 seconds (Royauté et al. 2020).

We conducted four latency trials at the same time in one of four arenas. We thoroughly cleaned each arena and artificial burrows with 70% ethanol wipes to avoid accumulation of chemical traces of conspecifics between trials.

### Antipredator Response

To measure responses to cues of potential predator presence, we collected excreta from three adult *E. maculariu*s, that were fed a diet of *G. sigillatus*. We froze the excreta and then finely ground and diluted it with deionized water (1 ml H_2_O : 5 mg of excreta). We applied this solution to 15 cm diameter filter paper disks with a 5 cm diameter central cutout that allows crickets to be left to rest unexposed to the predator cues (Royauté and Dochtermann 2017, Royauté et al. 2019). Each predator cue disk was allowed to dry for a minimum of 2 hours then stored at 4°C but was allowed to warm to room temperature before use in antipredator trials and discarded after a single use. We placed the predator cue disk at the bottom of a 15 cm diameter circular arena and kept the cricket for a minimum of 60 seconds under a 5 cm diameter cup in the nontreated central cutout. We then allowed the cricket to move freely for 220 seconds and estimated the distance travelled in cm using Ethovision XT.

We conducted four antipredator response trials at the same time in one of four arenas. We thoroughly cleaned each arena with 70% ethanol wipes to avoid accumulation of chemical traces of conspecifics between trials. We recorded the mass of each individual to the nearest 1 mg following the Antipredator Response assay.

Previous studies with this protocol have shown that crickets had heightened activity levels in the presence of this diluted gecko excreta compared to water controls (Royauté and Dochtermann 2017). Greater activity during the antipredator response assays was therefore interpreted as a greater responsiveness to predator cues.

### Data Analysis

To assess the effects of prolonged exposure to predators and predator stimuli we used mixed-effects models. We included sex, temperature (C), mass (mg), the number of days between maturity and testing date, and testing arena as fixed effects. We also included treatment, generation, and the interaction between them as fixed effects. If chronic exposure to predator cues resulted in a plastic response, we expected the treatments to differ between each other for adults of the F_0_ generation. If chronic exposure of parents produced a transgenerational effect, we expected the treatments to differ in the F_1_ generation.

For random effects, container ID was used for the F_0_ generation to account for non-independence among the 10 terraria in which subjects were reared. We also included parental pair ID of F_1_ individuals as a random effect. For the F_1_ generation, individuals were individually reared but all F_1_ individuals were given the same dummy container ID for analyses. Using unique container IDs for F_1_ individuals resulted in the within-container variation being improperly estimated. Similarly, F_0_ individuals were all assigned the same dummy pair ID due to their unknown relatedness. Using unique pair IDs for all F_0_ individuals again misestimated within-pair variation estimation. Consequently, our use of dummy values for F_0_ parents and F_1_ containers best accommodated the structure of our data while allowing for proper estimation of within-factor variances.

As a post-hoc analysis, differences in mass between generations and treatments were also estimated using mixed-effects models. We again included generation, treatment, their interaction, sex, temperature, number of days between maturity and testing date, and arena as fixed effects. Similar to mixed-effects models assessing behavioral differences, pair IDs and container IDs were also used for both generations in estimating differences in mass for treatment effects.

## Results

There were no effects of treatment or generation for anti-predator response or latency to emerge from shelter (Figure 2, Table 2). Male *G. sigillatus* moved ∼91 cm less than females (F_1,161.5_ = 5.66, p = 0.019; Table 2 & S1) during anti-predator trials.

**Table 2.** Model summary results for the effects of exposure (Treatment) on distance moved, time to emergence, and mass. Significant effects are in bold.

| Distance moved (cm) | Degrees of Freedom | F-value | p-value |
| --- | --- | --- | --- |
| Treatment | 1, 154.6 | 0.012 | 0.913 |
| Generation | 1, 7.2 | 3.228 | 0.114 |
| **Sex** | **1, 161.5** | **5.661** | **0.019** |
| Temperature | 1, 161.99 | 0.855 | 0.357 |
| Mass | 1, 163.24 | 0.382 | 0.537 |
| Days since maturation | 1, 143.96 | 0.009 | 0.926 |
| Arena | 3, 161.46 | 0.62 | 0.603 |
| Treatment × Generation | 1, 33.21 | 0.009 | 0.924 |
| Time to emergence (s) |  |  |  |
| Treatment | 1, 6.41 | 0.093 | 0.771 |
| Generation | 1, 6.29 | 0.441 | 0.53 |
| Sex | 1, 158.31 | 0.052 | 0.82 |
| Temperature | 1, 161.17 | 0 | 0.986 |
| Mass | 1, 163.1 | 0.761 | 0.384 |
| Days since maturation | 1, 130.65 | 0.005 | 0.947 |
| Arena | 3, 162.11 | 0.689 | 0.56 |
| Treatment × Generation | 1, 10.82 | 0.254 | 0.624 |
| Mass (mg) |  |  |  |
| **Treatment** | **1, 4.41** | **8.197** | **0.041** |
| **Generation** | **1, 2.7** | **14.793** | **0.037** |
| **Sex** | **1, 180.92** | **178.32** | **<0.01** |
| Temperature | 1, 172.95 | 1.081 | 0.3 |
| **Days since maturation** | **1, 122.13** | **3.959** | **0.049** |
| Arena | 3, 179.38 | 0.668 | 0.573 |
| **Treatment × Generation** | **1, 11.02** | **9.138** | **0.012** |

**Figure 2.**
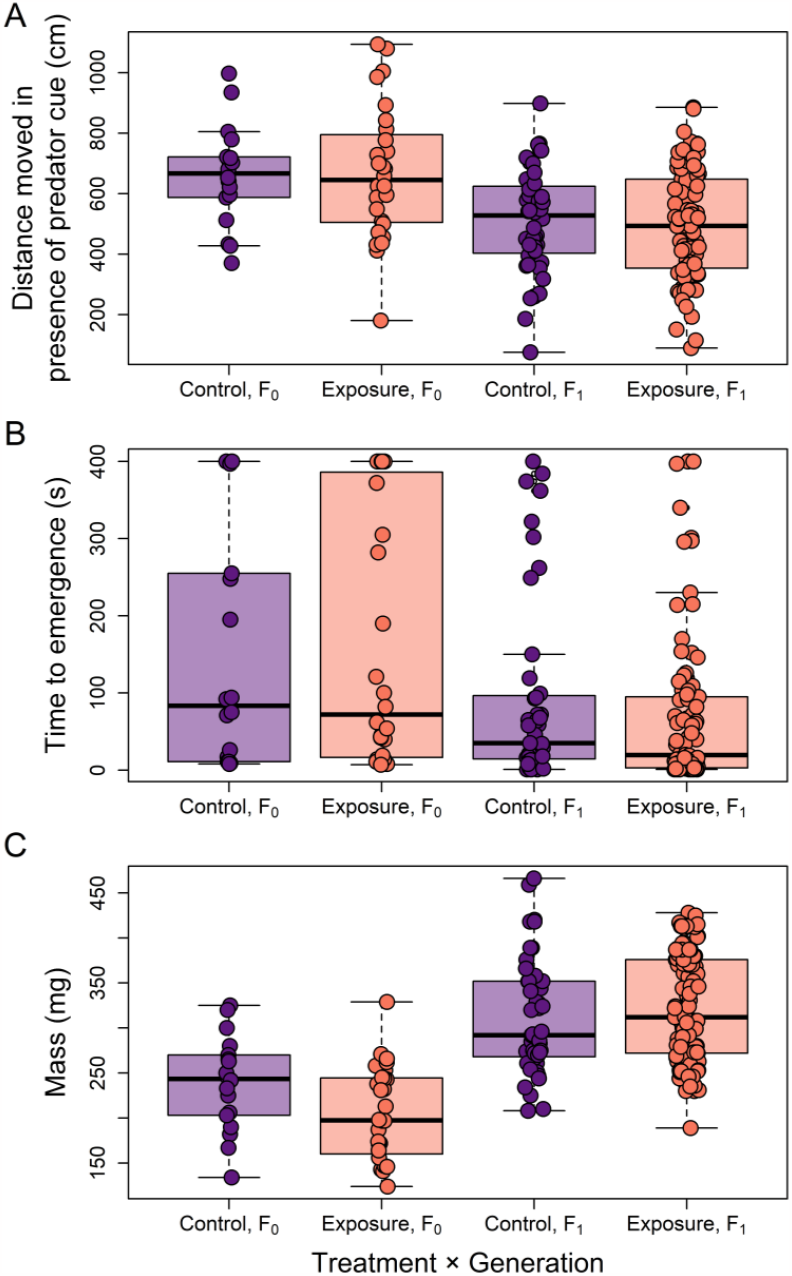
Differences in (A) distance traveled (cm) when exposed to cues of predator presence, (B) time to emergence from shelter (s), and (C) mass (mg) for control and treatment individuals by generation. Boxes indicate the lower and upper quartiles, horizontal lines within the boxes indicate the median, whiskers extend to 1.5 interquartile range of the box, and points indicate individual values. Differences by treatment, generation, and their interaction were not significant for distance moved or time to emergence (Table 2). All three were significant for mass (Table 2).

We found differences in mass due to treatment, generation, their interaction, sex, and days since maturation (Table 2). The F_1_ generation overall weighed ∼44 mg more than the F_0_ generation, regardless of treatment (Figure 2 & S1).

## Discussion

We failed to detect either a plastic behavioral response to chronic exposure or transgenerational effects of chronic exposure on behavior. Because we were able to detect both for mass (Figure 2, Table 2), we do not consider this to be due to insufficient statistical power. Instead, this failure to detect either suggests a lack, or loss, of a plastic response and a lack of transgenerational effects.

The failure to detect a plastic response is surprising because we have previously demonstrated that exposure to leopard geckos elicits behavioral plasticity in *G. sigillatus* (Bucklaew and Dochtermann 2020). Specifically, Bucklaew and Dochtermann (2020) showed that when *G. sigillatus* are directly confronted with a leopard gecko predator they subsequently increase their activity in an open field behavioral assay—even without predator cures being present—and delay emergence from shelter. A major difference between these studies is that here F_0_ individuals were only tested once at maturation to prevent repeated exposure to the cue for the control groups. Given that we only measured behaviors at maturation and that acute exposure has been shown to elicit a response, subjects may have had an initial response to the presence of the leopard geckos and, as exposure continued, behavioral responses habituated back to the same level as the control group.

The absence of detectable transgenerational effects is similarly surprising. Storm and Lima (2010) demonstrated that other Gryllidae crickets when, similarly, chronically exposed to predators and their cues had offspring that were more responsive to predation threat. A chief difference between the current study and that of Storm and Lima (2010) is that, here, crickets were not gravid during the period of exposure. Three possible explanations for our results and their inconsistency with those of others are therefore: i) crickets here habituated to cues of predator presence due to the lack of direct interaction and attacks; ii) cues alone were insufficient to induce transgenerational effects; or iii) differences in responses to vertebrate versus invertebrate predators. We cannot currently distinguish among these alternatives.

Differences in mass at maturity might indicate the effects of stress caused by the direct exposure to cues of predator presence during development. The physiological effects of stress on development are well document (DeVries et al. 1997, Mishra et al. 2011, Kriengwatana et al. 2013, Royauté et al. 2019) and significantly lower body masses could be an indicator of a stressful developmental environment caused by the presence of live leopard geckos (Kriengwatana et al. 2013). Evidence of F_0_ treatment individuals initially responding to the presence of the leopard geckos would then be suggested by the experimental group having significantly lower masses at maturation compared to control individuals (Figure 2C). This potential for developmental plasticity (sensu West-Eberhard 2003) on mass during development did not, however, produce transgenerational effects in the same direction on offspring mass as F_1_ treatment individuals weighed significantly more than the previous generation (Figure 2C). Significantly greater masses overall of the F_1_ generation compared to the F_0_ generation is most likely due to the increased quality of rearing conditions as the F_1_ generation was reared in individual containers and without the presence of a live predator. However, the effects of prolonged exposure to predator cues during development needs to be further investigated as the findings of differences in mass were not part of our *a priori* questions and were instead discovered during post-hoc data exploration.

Our results, combined with recent findings, suggest that after initial responses to predator presence prey will revert behaviors back to baseline levels of responsiveness if not reinforced by direct consequences of predator interaction. For example, similar patterns were identified by Pilakouta and Alonzo (2014), where the authors observed that female *Xiphophorus helleri* (green swordtail fish) changed their mate preference from males with longer swords to males with shorter swords when in the presence of predators as compared to a control group. This altered mate choice only lasted for 24 hours after predator exposure and female preference shifted back towards males with longer swords, consistent with control individual preference. This lends further support to the argument that behavioral responses can be influenced by predator exposure but, without continued exposure, these temporary behavioral changes can revert to the normal levels observed before predator exposure.

The lack of apparent behavioral response to chronic exposure of predator cues raises several questions. First, if, as our results and the behavior of the species in other contexts suggests, habituation resets behaviors back to naïve levels, at what point in exposure duration does the habituation occur? Second, since a lack of differences in behavioral responses of treatment groups contradicts responses to extreme or acute exposure examples found in the literature (Storm and Lima 2010, Bucklaew and Dochtermann 2020), the effects of prolonged exposure to predation stimuli remains unclear.

## Acknowledgements

The authors thank M. Anderson-Berdal, E. Gillam, J. Harmon, C. Hickey, and R. Royaute for helpful guidance and advice throughout. We also thank Brady Klock, Ishan Joshi, Hannah Lambert, Jenna LaCoursiere and Maddi Rick for assistance in conducting behavioral trials and in rearing and care of the crickets. Finally, we thank S. Sakaluk for sharing his lines of crickets. This work was supported by U.S. NSF IOS grant 1557951 to NAD.

## Supplementary Information

**Table S1.** Linear model coefficients for each of the three traits. The intercept was calculated for F_0_ females in the control treatment when measured in arena one (of four) and at the average temperature behavioral assays were conducted. Coefficients for factors are differences between the intercept/reference and other levels.

| Distance moved (cm) | Estimate | Std. Error | df | t value | Pr(>\|t\|) |
| --- | --- | --- | --- | --- | --- |
| (Intercept) | 758.9108 | 126.7227 | 34.66 | 5.989 | <0.001 |
| Treatment (exposure) | -6.0158 | 54.9903 | 154.5986 | -0.109 | 0.913 |
| Generation (F<sub>1</sub>) | -161.2565 | 102.2673 | 9.3274 | -1.577 | 0.148 |
| Sex (males) | -91.1479 | 38.3094 | 161.5035 | -2.379 | 0.019 |
| Temperature | 36.3699 | 39.3361 | 161.9858 | 0.925 | 0.357 |
| Mass | -0.2094 | 0.3387 | 163.2361 | -0.618 | 0.537 |
| Days since maturation | 0.7742 | 8.3681 | 143.962 | 0.093 | 0.926 |
| Arena 2 | 25.2161 | 36.7046 | 161.1853 | 0.687 | 0.493 |
| Arena 3 | 27.4268 | 38.1919 | 162.0099 | 0.718 | 0.474 |
| Arena 4 | -21.4955 | 41.2316 | 162.5632 | -0.521 | 0.603 |
| Treatment × Generation | 7.8025 | 81.2544 | 33.2127 | 0.096 | 0.924 |

| Time to emergence (s) | Estimate | Std. Error | df | t value | Pr(>\|t\|) |
| --- | --- | --- | --- | --- | --- |
| (Intercept) | 199.1306 | 93.4015 | 34.7282 | 2.132 | 0.040 |
| Treatment (exposure) | 20.7217 | 68.0715 | 6.4056 | 0.304 | 0.771 |
| Generation (F<sub>1</sub>) | -48.4579 | 112.5322 | 7.1701 | -0.431 | 0.679 |
| Sex (males) | 5.7423 | 25.2107 | 158.3139 | 0.228 | 0.820 |
| Temperature | -0.4553 | 26.4923 | 161.1676 | -0.017 | 0.986 |
| Mass | -0.1946 | 0.2231 | 163.0973 | -0.872 | 0.384 |
| Days since maturation | -0.3759 | 5.5867 | 130.6472 | -0.067 | 0.947 |
| Arena 2 | -5.2468 | 24.1554 | 161.8785 | -0.217 | 0.828 |
| Arena 3 | 17.9235 | 24.7918 | 159.6055 | 0.723 | 0.471 |
| Arena 4 | -20.7202 | 26.8232 | 164.4857 | -0.772 | 0.441 |
| Treatment × Generation | -39.6608 | 78.627 | 10.8202 | -0.504 | 0.624 |

| Mass (mg) | Estimate | Std. Error | df | t value | Pr(>\|t\|) |
| --- | --- | --- | --- | --- | --- |
| (Intercept) | 283.328 | 17.476 | 7.006 | 16.213 | <0.001 |
| Treatment (exposure) | -38.466 | 13.436 | 4.405 | -2.863 | 0.041 |
| Generation (F<sub>1</sub>) | 44.608 | 22.382 | 4.16 | 1.993 | 0.114 |
| Sex (males) | -79.894 | 5.983 | 180.92 | -13.354 | <0.001 |
| Temperature | 8.968 | 8.624 | 172.951 | 1.04 | 0.300 |
| Days since maturation | 3.343 | 1.68 | 122.125 | 1.99 | 0.049 |
| Arena 2 | 3.769 | 7.901 | 179.166 | 0.477 | 0.634 |
| Arena 3 | 2.384 | 8.263 | 181.225 | 0.289 | 0.773 |
| Arena 4 | 12.068 | 8.796 | 183.678 | 1.372 | 0.172 |
| Treatment × Generation | 54.261 | 17.95 | 11.023 | 3.023 | 0.012 |

## Notes

### Competing Interest Statement

The authors have declared no competing interest.

## Literature Cited

Binz, H., R. Bucher, M. H. Entling, and F. Menzel. 2014. Knowing the risk: crickets distinguish between spider predators of different size and commonness. Ethology 120:99–110.

Bucklaew, A., and N. Dochtermann. 2020. The effects of exposure to predators on personality and plasticity. Ethology.

Committee, A. A. A. C. 2023. Guidelines for the ethical treatment of nonhuman animals in behavioural research and teaching. Animal Behaviour 195:I–XI.

Dalos, J., R. Royauté, A. Hedrick, and N. A. Dochtermann. 2022. Phylogenetic conservation of behavioural variation and behavioural syndromes. Journal of Evolutionary Biology 35:311–321.

DeVries, A. C., J. M. Gerber, H. N. Richardson, C. A. Moffatt, G. E. Demas, S. E. Taymans, and R. J. Nelson. 1997. Stress affects corticosteroid and immunoglobulin concentrations in male house mice (Mus musculus) and prairie voles (Microtus ochrogaster). Comparative Biochemistry and Physiology Part A: Physiology 118:655–663.

Dochtermann, N. A., T. Schwab, M. A. Berdal, J. Dalos, and R. Royaute. 2019. The heritability of behaviour: a meta-analysis. Journal of Heredity 110:403–410.

Eklöv, P., and T. VanKooten. 2001. Facilitation among piscivorous predators: effects of prey habitat use. Ecology 82:2486–2494.

Ivy, T. M., C. B. Weddle, and S. K. Sakaluk. 2005. Females use self-referent cues to avoid mating with previous mates. Proceedings Of The Royal Society B-Biological Sciences 272:2475–2478.

Kalb, N., and C. Randler. 2019. Behavioral responses to conspecific mobbing calls are predator-specific in great tits (Parus major). Ecology and Evolution 9:9207–9213.

Kikuchi, D. W., W. L. Allen, K. Arbuckle, T. G. Aubier, E. S. Briolat, E. R. Burdfield-Steel, K. L. Cheney, K. Daňková, M. Elias, L. Hämäläinen, M. E. Herberstein, T. J. Hossie, M. Joron, K. Kunte, B. C. Leavell, C. Lindstedt, U. Lorioux-Chevalier, M. McClure, C. F. McLellan, I. Medina, V. Nawge, E. Páez, A. Pal, S. Pekár, O. Penacchio, J. Raška, T. Reader, B. Rojas, K. H. Rönkä, D. C. Rößler, C. Rowe, H. M. Rowland, A. Roy, K. A. Schaal, T. N. Sherratt, J. Skelhorn, H. R. Smart, T. Stankowich, A. M. Stefan, K. Summers, C. H. Taylor, R. Thorogood, K. Umbers, A. E. Winters, J. Yeager, and A. Exnerová. 2023. The evolution and ecology of multiple antipredator defences. Journal of Evolutionary Biology 36:975–991.

Kortet, R., A. Vainikka, M. Janhunen, J. Piironen, and P. Hyvarinen. 2014. Behavioral variation shows heritability in juvenile brown trout Salmo trutta. Behavioral Ecology and Sociobiology 68:927–934.

Kriengwatana, B., H. Wada, A. Macmillan, and S. A. MacDougall-Shackleton. 2013. Juvenile nutritional stress affects growth rate, adult organ mass, and innate immune function in zebra finches (Taeniopygia guttata). Physiological and Biochemical Zoology 86:769–781.

Makin, D. F., S. Chamaillé-Jammes, and A. M. Shrader. 2017. Herbivores employ a suite of antipredator behaviours to minimize risk from ambush and cursorial predators. Animal Behaviour 127:225–231.

Marshall, D. J., and T. Uller. 2007. When is a maternal effect adaptive? Oikos 116:1957–1963.

Miller, J. R., J. M. Ament, and O. J. Schmitz. 2014. Fear on the move: predator hunting mode predicts variation in prey mortality and plasticity in prey spatial response. Journal of Animal Ecology 83:214–222.

Mishra, S., D. M. Logue, I. O. Abiola, and W. H. Cade. 2011. Developmental environment affects risk-acceptance in the hissing cockroach, Gromphadorhina portentosa. Journal of Comparative Psychology 125:40.

Niemela, P. T., N. DiRienzo, and A. V. Hedrick. 2012. Predator-induced changes in the boldness of naive field crickets, Gryllus integer, depends on behavioural type. Animal Behaviour 84:129–135.

Palmer, M. S., and C. Packer. 2021. Reactive anti-predator behavioral strategy shaped by predator characteristics. Plos One 16:e0256147.

Pilakouta, N., and S. H. Alonzo. 2014. Predator exposure leads to a short-term reversal in female mate preferences in the green swordtail, Xiphophorus helleri. Behavioral Ecology 25:306–312.

Preisser, E. L., J. L. Orrock, and O. J. Schmitz. 2007. Predator hunting mode and habitat domain alter nonconsumptive effects in predator–prey interactions. Ecology 88:2744–2751.

Royauté, R., and N. A. Dochtermann. 2017. When the mean no longer matters: developmental diet affects behavioral variation but not population averages in the house cricket (Acheta domesticus). Behavioral Ecology 28:337–345.

Royauté, R., C. Garrison, J. Dalos, M. A. Berdal, and N. A. Dochtermann. 2019. Current energy state interacts with the developmental environment to influence behavioural plasticity. Animal behaviour 148:39–51.

Royauté, R., K. Greenlee, M. Baldwin, and N. A. Dochtermann. 2015. Behaviour, metabolism and size: phenotypic modularity or integration in Acheta domesticus? Animal Behaviour 110:163–169.

Royauté, R., A. Hedrick, and N. A. Dochtermann. 2020. Behavioural syndromes shape evolutionary trajectories via conserved genetic architecture. Proceedings of the Royal Society B 287:20200183.

Snell-Rood, E. C. 2013. An overview of the evolutionary causes and consequences of behavioural plasticity. Animal Behaviour 85:1004–1011.

Storm, J. J., and S. L. Lima. 2010. Mothers Forewarn Offspring about Predators: A Transgenerational Maternal Effect on Behavior. American Naturalist 175:382–390.

Thompson, R. F. 2009. Habituation: a history. Neurobiology of learning and memory 92:127–134.

Van Buskirk, J. 2001. Specific induced responses to different predator species in anuran larvae. Journal of Evolutionary Biology 14:482–489.

West-Eberhard, M. J. 2003. Developmental Plasticity and Evolution. Oxford University Press, New York.

